# A development guide for evaluating the maximum yield potential stage in barley

**DOI:** 10.1101/2021.02.02.429383

**Authors:** Venkatasubbu Thirulogachandar, Thorsten Schnurbusch

## Abstract

Determining the grain yield potential contributed by grain number is a step towards advancing cereal crops’ yield. To achieve this aim, it is pivotal to recognize the maximum yield potential (MYP) of the crop. In barley (*Hordeum vulgare* L.), the MYP is defined as the maximum spikelet primordia number of a spike. Previous barley studies often assumed the awn primordium (AP) stage as the MYP stage regardless of genotypes and growth conditions. From our spikelet-tracking experiments using the two-rowed cultivar Bowman, we found that the MYP stage can be different from the AP stage. Importantly, we find that the occurrence of inflorescence meristem (IM) deformation and its loss of activity coincided with the MYP stage, indicating the end of further spikelet initiation. Thus, we recommend validating the barley MYP stage with the IM’s shape and propose this approach (named Spikelet Stop) for MYP staging. Following this approach, we compared the MYP stage and the MYP in 27 two- and six-rowed barley accessions grown in the greenhouse and field. Our results reveal that the MYP stage can be reached at various developmental stages, which majorly depend on the genotype and growth conditions. Furthermore, we found that two-rowed barleys’ MYP and the duration reaching the MYP stage may determine their yield potential. Based on our findings, we suggest key steps for the identification of the MYP in barley that can also be applied in a related crop such as wheat.

**Highlight:** We show that the maximum yield potential stage in barley can be different from the awn primordium stage as proposed in earlier studies and it varies depending on the genotype and growth conditions. We suggest key steps to identify maximum yield potential in barley that might apply to related cereals.

## Introduction

Grain yield potential is the key to augmenting our crops’ yield that warrants the subsistence of humanity. In the small-grain cereals, such as wheat (*Triticum* spp.) and barley (*Hordeum vulgare*, L), grain number (GN) per unit area shows a significant influence on the final grain yield (Prystupa et al., 2004; Arisnabarreta and Miralles, 2006b; Peltonen-Sainio et al., 2007; Ugarte et al., 2007; Serrago et al., 2013; Arisnabarreta and Miralles, 2015; García et al., 2015; Sakuma and Schnurbusch, 2020). Wheat and barley belong to the tribe Triticeae and prototypically possess an unbranched inflorescence known as the spike. The spike of these crops follows a distichous phyllotactic arrangement of spikelets (basic floral units of the spike) on its rachis (spike axis), whereas architecturally, the spike apex (inflorescence meristem) and spikelets are different in these crops. In wheat, the spike is determinate, i.e., it culminates spikelet initiation by forming a terminal spikelet; however, the barley spike is indeterminate and produces spikelets indefinitely without a terminal spikelet. Furthermore, wheat develops a single spikelet on every rachis node, and its spikelet produces florets (reduced flowers) indeterminately. On the contrary, barley forms three spikelets at every node, while its spikelets are determinate and bear a single floret (Bonnett, 1935; Bonnett, 1936; Whipple, 2017; Bommert and Whipple, 2018; McKim et al., 2018; Gauley and Boden, 2019; Koppolu and Schnurbusch, 2019).

In wheat and barley, GN is majorly determined by the development and survival of the florets and spikelets (Arisnabarreta and Miralles, 2008a; Bancal, 2008, 2009; Gonzalez et al., 2011; Ferrante et al., 2012; Ferrante et al., 2013; Arisnabarreta and Miralles, 2015; Digel et al., 2015; Sakuma et al., 2019). To fathom the surviving portion of the florets/spikelets, it is necessary to identify the maximum number of florets/spikelets, also popularly known as ‘maximum yield potential’ (MYP). In wheat, MYP is ascertained by trailing the floret number of spikelets (indeterminate) located at various spike positions (basal, central, and apical) until the spikelets cease floret initiation (Gallagher, 1979; Ferrante et al., 2013; Guo and Schnurbusch, 2015; Ferrante et al., 2020). To define the MYP in barley, the spikelet/floret number per spike needs to be followed up until the stage at which the indeterminate barley spike ceases spikelet initiation (Gallagher et al., 1976; Kirby, 1977; Appleyard et al., 1982; Arisnabarreta and Miralles, 2006a; Arduini et al., 2010). Studies above implied that the MYP must be discerned by following the indeterminate floral structure (spikelet or spike) until it stops its activity. By enumerating the spikelet primordia number from sowing until anthesis, Appleyard et al. (1982) conducted a study using three barley genotypes, including their crosses. They identified the maximum number of spikelets after spikelet initiation had stopped (Fig.1 in Appleyard et al., 1982). The authors found that the number of spikelets counted at the MYP stage was close to the spikelet number at the awn primordium (AP) stage (Kirby and Appleyard, 1984) since the average difference was only 0.52 primordium. It seems astonishing; but in fact, all subsequent studies presumed that barley’s MYP is around the AP stage irrespective of the genotypes and the growth conditions used in such experiments (Kirby and Appleyard, 1984; del Moral et al., 1991; Miralles et al., 2000; Arisnabarreta and Miralles, 2006a; Arisnabarreta and Miralles, 2010; Alqudah and Schnurbusch, 2014; Arisnabarreta and Miralles, 2015). This initial assumption appears simplistic and premature; and thus, rather requires more experimental verification to better understand the relevance of the AP stage corresponding to the MYP using various barley accessions and growth conditions.

**Figure 1:**
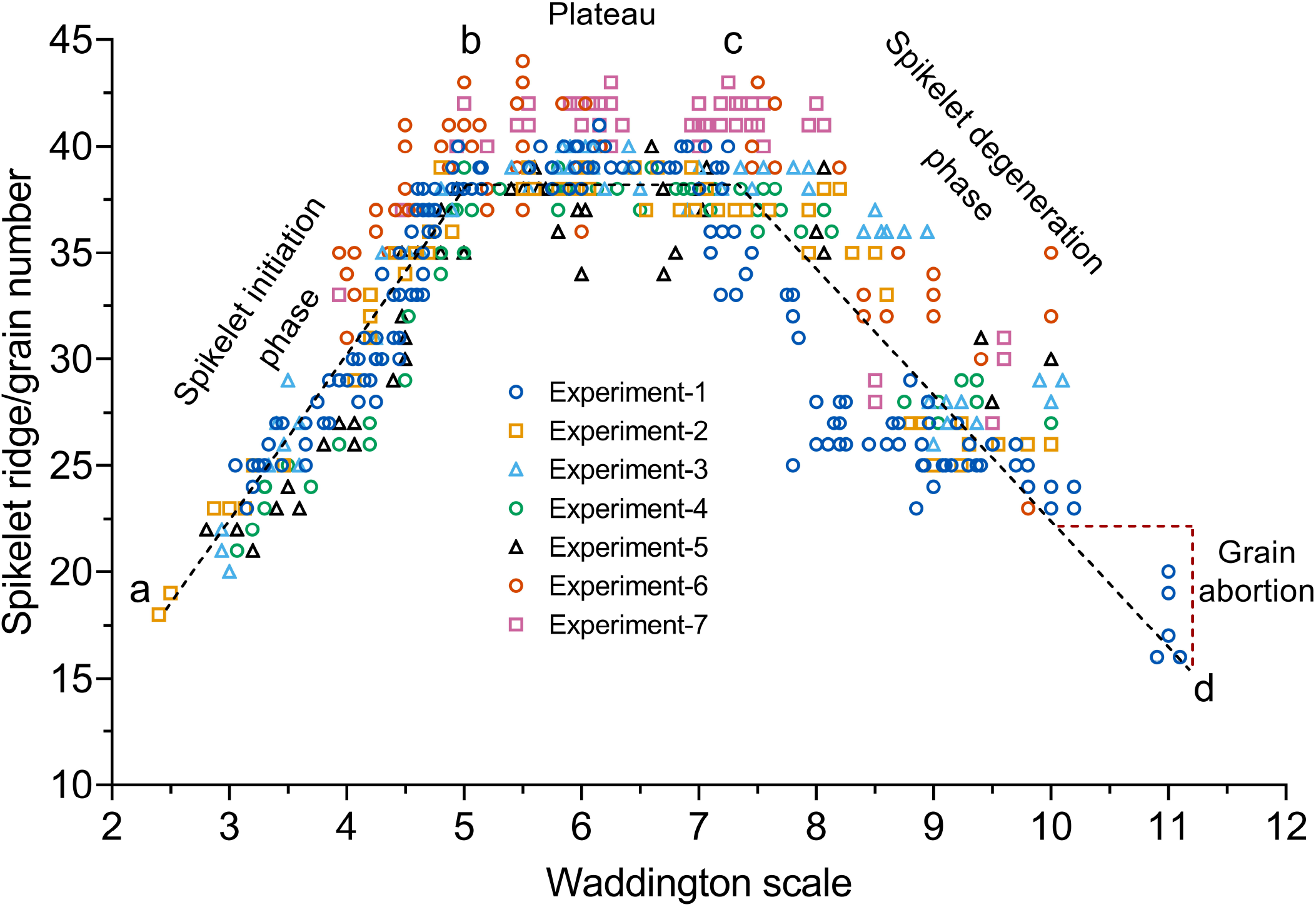
The spikelet/floret initiation and growth pattern of barley. Spikelet initiation and growth pattern of a barley cv. Bowman analyzed in seven different experiments is shown. It shows that the pattern has three phases - the first phase (from point ‘a to b’) indicates the increase of spikelet ridge number during the spikelet initiation phase. The second phase (from ‘b to c’) is a plateau during which the number of spikelet ridges did not change, and the last phase (from ‘c to d’) displays the spikelet degenerative phase and grain abortion phase. The inflection point ‘b’ marks the end of the spikelet initiation phase or the stage of maximum yield potential (MYP). In contrast, point ‘c’ is the initiation of the spikelet abortion phase. Each stage may have data from one to nine different plants.

In the present study, we, therefore, analyzed the MYP stage on the main culm of a two-rowed cv. Bowman grew in seven different experiments (variable growth conditions and pot sizes). In these experiments, we identified the MYP stage by following the spikelet number per spike until the cessation of spikelet initiation. Among the experiments, in only two instances, Bowman reached the MYP stage close to the AP stage, indicating that the MYP stage can be different from the previously proposed AP stage (Appleyard et al., 1982). In fact, we demonstrate that morphological changes of the inflorescence meristem (IM) during spikelet initiation and growth are highly linked with reaching the MYP stage and the cessation of spikelet initiation. This method for identifying the MYP stage was termed ‘Spikelet Stop’ (SS) as it axiomatically defines ‘spikelets at the end of spikelet initiation.’ Furthermore, by applying the SS method for the identification of the MYP stage and the MYP, we tracked the developmental progression following the Waddington scale (Waddington et al., 1983) and the number of spikelets on the main culm spike in a panel of two-(17) and six-rowed (10) genotypes grown in the greenhouse and field. Our findings from this panel clarify unambiguously that the MYP stage can occur at various developmental stages and that the MYP depends on genotype and growth conditions. Interestingly, we also observed that the timing/duration and the stage reaching MYP in two-rowed types’ might determine the grain number of the main culm spike. Finally, this study provides an appropriate methodology for the identification of the MYP in barley spikes.

## Materials and Methods

### Plant materials and growth conditions

We conducted one greenhouse and one field experiment using a panel of twenty-seven barley accessions chosen from a worldwide collection (Alqudah et al., 2014). The field study was conducted at the Leibniz Institute of Plant Genetics and Crop Plant Research, Gatersleben, Germany (51° 49′ 23″ N, 11° 17′ 13″ E, altitude 112 m) from March to August 2017 and greenhouse from July to December 2017. For both the experiments, we followed similar sowing and pre-treatment protocols. Grains were sown on 96 well trays and grown under greenhouse conditions (photoperiod, 16h/8h, light/dark; temperature, 20° C/16° C, light/dark) for two weeks. Then, the grown plants were vernalized at 4° C for four weeks and then given hardening in the greenhouse for a week. Following the acclimatization, plants were directly transplanted in the silty loam soil for the field experiment; however, in the greenhouse experiment (photoperiod, 16h/8h, light/dark; temperature, 20° C/16° C, light/dark), plants were potted in a 9 cm pot (9 × 9 cm, diameter × height). For the field experiment, we followed a single plot per genotype design, in which each plot had eight rows with a 15 cm distance between the rows. The rows are 0.8 m long, wherein we sowed five plants per row. Details of the selected barley panel and their field growth conditions were given in tables S1 and S2, respectively.

The series of experiments on cultivar Bowman were performed in the greenhouse and climate chamber. The growth conditions and different pot sizes of the experiments are given in table S3. In all the greenhouse and climate chamber experiments, plants were grown in pots that contain two parts of autoclaved compost, two parts of ‘Rotes Substrat’ (Klasmann-Deilmann GmbH, Germany), and one part of sand. We followed the standard practices for irrigation, fertilization, and control of pests and diseases in these experiments.

### Methods of phenotyping the traits

All traits were measured only from the main culm of a barley plant due to its higher phenotypic stability across growth conditions and its major contribution to the final grain yield (Cottrell et al., 1985; Elhani et al., 2007). In all Bowman spikelet-tracking experiments, plants were randomly selected, and spikes were dissected out almost every alternate day. In the spikelet-tracking experiments with the 27 accessions, random plants were dissected every two to three days or more, depending on their developmental rate. Dissection of spikes was performed according to the methods described in Kirby and Appleyard, 1984. Different spike developmental stages were identified by following the description provided earlier (Waddington et al., 1983; Kirby and Appleyard, 1984). For every stage of a spike, the decimal code suggested by Waddington et al., 1983 was given following the letter ‘W’ (Waddington). A decimal code was assigned to a spike when the specific stage was found in a minimum of three or a maximum of four consecutive nodes of a spike (Waddington et al., 1983; Kirby and Appleyard, 1984). These nodes are always the most developed nodes of a spike (Kirby and Appleyard, 1984) and may be found close to the spike base, two to three nodes above the spike base, or in the center of a spike.

For the potential spikelet number (PSN) of a spike, differentiated spikelets and undifferentiated spikelet ridges (usually found at the base and tip of a spike) were counted (Appleyard et al., 1982). Generally, barley forms three spikelets at every rachis node (Bonnett, 1935; Bonnett, 1966; Komatsuda et al., 2007; Koppolu et al., 2013), so the number of undifferentiated spikelet ridges were multiplied by three. The number of ridges developed on a spike was considered as spikelet ridge number (SRN), and the final spikelet number (SN) and grain number (GN) of a spike were enumerated after physiological maturity. Growing degree days (GDD) or thermal time was calculated from the average maximum and minimum air temperature of a day subtracted by the base temperature (McMaster and Wilhelm, 1997). We considered 0° C as the base temperature for barley (Gallagher et al., 1976). GDDs and PSN/SRN were taken from three plants, while SN and GN were from six plants.

### Data analysis

The data were analyzed using the Prism software, version 8.4.2 (GraphPad Software, LLC), and outliers were detected by the ‘ROUT’ method (Motulsky and Brown, 2006). Mean value comparison of different traits was made with either the multiple Student’s t-tests or paired Student’s t-test (parametric). The false discovery rate approach of the two-stage linear step-up procedure of Benjamini, Kreier, and Yekutieli (Q=5%) (Benjamini et al., 2006) was used to calculate the significance of Student’s t-tests. For traits like the Waddington scale and Growing Degree Days at MYP, a two-way ANOVA with Tukey’s multiple comparison test (alpha=5%) was used to identify the mean values’ significance. For the spikelet ridge number (SRN), a one-way ANOVA with Tukey’s multiple comparison test (alpha=5%) was used to identify the mean values’ significance. All the replicates of a genotype were analyzed individually and without assuming a consistent standard deviation. Linear regression was done using the appropriate dependent (Y values) and independent (X values) traits. The 95% confidence intervals were identified for every linear regression and plotted as confidence bands along with the ‘goodness of fit’ line.

## Results

### The spikelet/floret initiation and growth pattern of barley

We conducted seven experiments (variable pot sizes and growth conditions) using the two-rowed barley cv. Bowman, in which spikelet ridges were counted from the glume primordium stage (Waddington, W2.5) until pollination (W10.0) (Waddington et al., 1983). Additionally, we enumerated the final grain number (only in Experiment 1) of main culm spikes after harvest (considered as W11.0) and plotted the values together with the spikelet ridge number (SRN) (Fig.1
). Here, we enumerated every rachis node with differentiated and undifferentiated spikelet primordia and regarded them as ‘spikelet ridges.’ The charted values of spikelet ridges and grains exhibited a pattern that shows three distinct phases of spikelet initiation and growth. The entire pattern starts from point ‘a’ and ends in point ‘d’ (Fig. 1). In addition to these two stages (start and end), there are two inflection points, ‘b’ and ‘c.’ The phase between moment ‘a’ to ‘b’ shows a steady increase of SRN produced from ~W2.5 to ~W5.0 (Fig.1
) and represents the spikelet/floret initiation phase (Gallagher, 1979; Appleyard et al., 1982). Following the initiation phase, the period between the inflection points ‘b and c’ is a plateau (Appleyard et al., 1982), during which the SRN is generally unchanged (Fig. 1). While new spikelet initiation has stopped, spikelets and spikelet ridges formed during the initiation phase continue to grow and differentiate during the plateau phase. The subsequent final phase starts at point ‘c’ and ends at point ‘d’, displaying the gradual decrease of SRN due to spikelet abortion; thus, it is considered the spikelet degeneration phase (Fig. 1). Notably, at the first inflection point (‘b’), a spike bears the maximum SRN that is already equivalent to the MYP, denoting the end of the spikelet initiation phase. Afterward, the SRN remains similar until the second inflection point (‘c’). At the same time, immediately after this stage (plateau), the spikelet/floret abortion process is initiated, which continues until (or around) the pollination stage (W10.0) (Waddington et al., 1983; Kirby and Appleyard, 1984) (Fig. 1). After pollination, there can still be a reduction in the spikelet/grain number; but here, it is often due to the deficiency of grain setting that leads to grain abortion (Gallagher, 1979) (Fig. 1). Thus, from our results, it is evident that barley spikelet/floret initiation and growth follows a typical pattern that was reported previously (Gallagher, 1979; Appleyard et al., 1982), and it includes a spikelet generation phase followed by a period of no further spikelet initiation (plateau), which in turn is trailed by the spikelet degeneration phase.

### Spikelet initiation arrest marks the MYP stage, and it can vary depending on growth conditions

The barley spike belongs to the indeterminate type of inflorescences that develops spikelet primordia in acropetal succession without forming a terminal spikelet (Bonnett, 1935; Bonnett, 1966). In Fig.2, we display the young Bowman spikes (immature inflorescences) (from experiment 7) of stages W4.0 to W7.0, showing the pattern of spikelet ridge development. To this end, we scored the carpel development of the corresponding stages (from W4.5 to W7.0) in Fig. S1. In W4.0, the total number of spikelet ridges was 33 (Fig. 2A), which got increased to 37 at W4.5 (Fig. 2B), and then escalated to 42 in W5.0 (Fig. 2C). Notably, after W5.0, the SRN was not drastically changed, at least four more consecutive stages, i.e., until W7.0 (Fig. 2D-G). Here, the culmination of SRN at W5.0 indicated the arrest of spikelet initiation as well as the loss of IM activity. The gradual loss of the meristematic potential of the IM was evident from the images of spike apices from stage W4.0 to W7.0 (Fig.3). At W4.0, the IM had a rounded tip (Fig. 3A) with a slightly reduced size at W4.5 (Fig. 3B). Intriguingly, the IM lost its rounded tip at ~W5.0 and modified it to an oblique shape (Fig. 3C). Subsequently, the IM never regained its smooth shape after W4.5 (Fig. 3B); instead, it became tapered in the following stages (Fig. 3D-G). The structural alteration of the IM (Fig. 3C) and the cessation of spikelet ridge formation (Fig. 2C) at ~W5.0 indicated that spikelet initiation got arrested at this stage; and thus, stage W5.0 specified the MYP stage of cv. Bowman. Because the MYP stages indicate the stage at which spikelet initiation stops, we termed the method for identifying the MYP stage by definition “Spikelet Stop” (SS). We also verified the MYP stage of experiment 7 by plotting its values and spotted the first inflection point of the spikelet initiation curve at W5.0 (Fig.4A). Additionally, we compared the SRN between the AP stage and MYP stage (identified by the SS method) from all our Bowman experiments. We found an elevation of SRN by three to eight ridges (at the defined MYP stage across experiments), again indicating that the MYP stage can be different from the AP stage (Fig. 4B). Only in experiment 1, the MYP stage (~W4.9) was close to the AP stage (W4.5) (Fig. S2A); however, in this experiment also the SRN is significantly higher than the AP stage (Fig. 4B). Thus, our spikelet-tracking studies using the cultivar Bowman unambiguously showed that the spikelet initiation arrest marks the MYP stage, which could be different from the AP stage (W4.5), and that the MYP stage varies depending on growth conditions (Fig, S2C).

**Figure 2:**
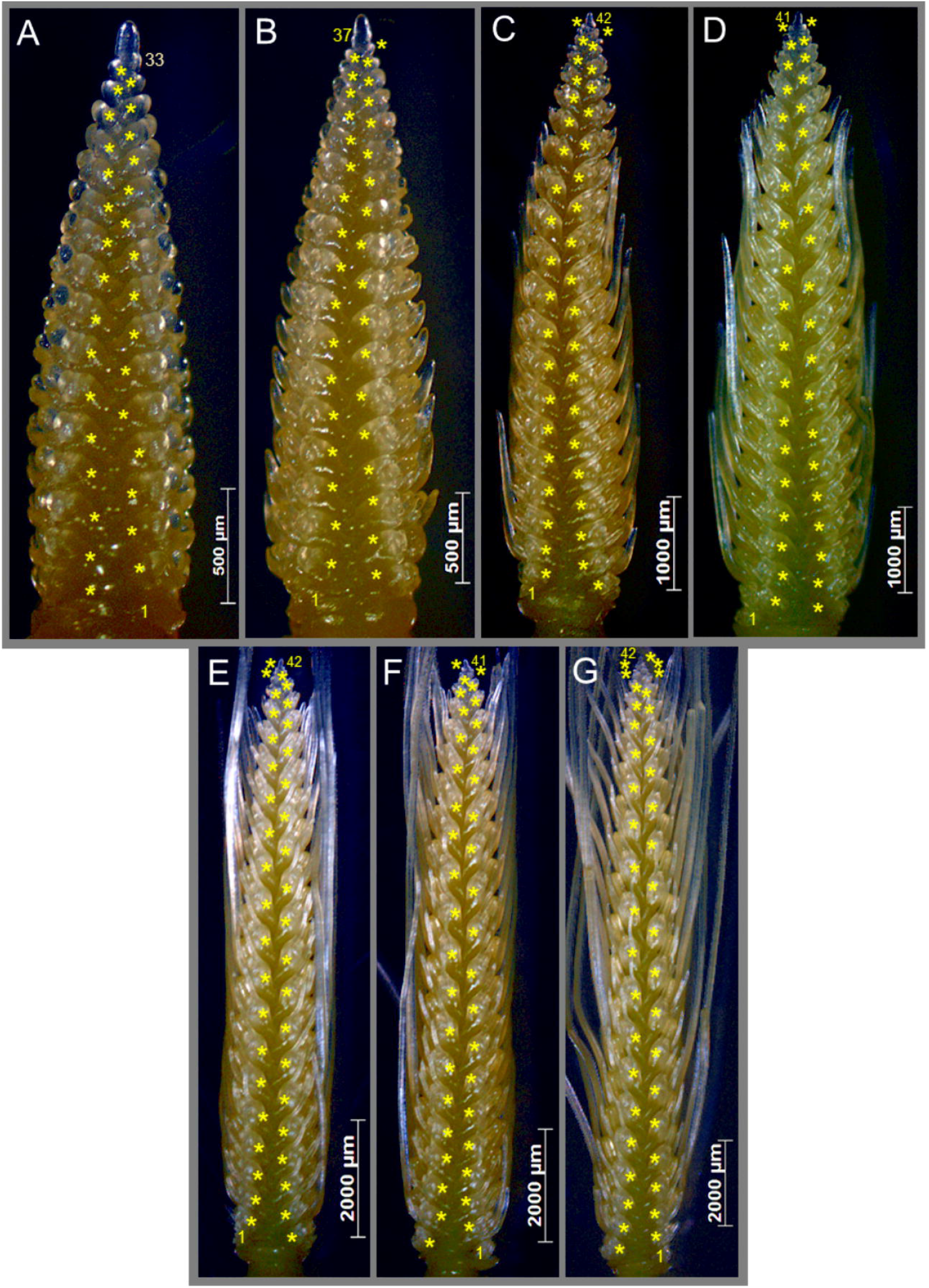
Arrest of spikelet initiation marks the maximum yield potential (MYP) stage in barley spikes. A representative spike of stages from W4.0 to W7.0 and their number of spikelet ridges are shown (A-G) from experiment 7. The number of spikelet ridges steadily increased from W4.0 to WS.O. There were 33 ridges at W4.0 (A) and elevated to 37 in W4.5 or awn primordium (AP) stage (B). The ridge number increased after the AP stage to 42 in WS.O (C). After stage WS.O, the number of ridges was not drastically elevated until W7.0 (D-G). These data clearly demonstrate that spikelet initiation stopped at WS.O. From these images, it thus becomes evident that the MYP stage was at WS.O and not at W4.5 (AP). W-Waddington scale.

**Figure 3:**
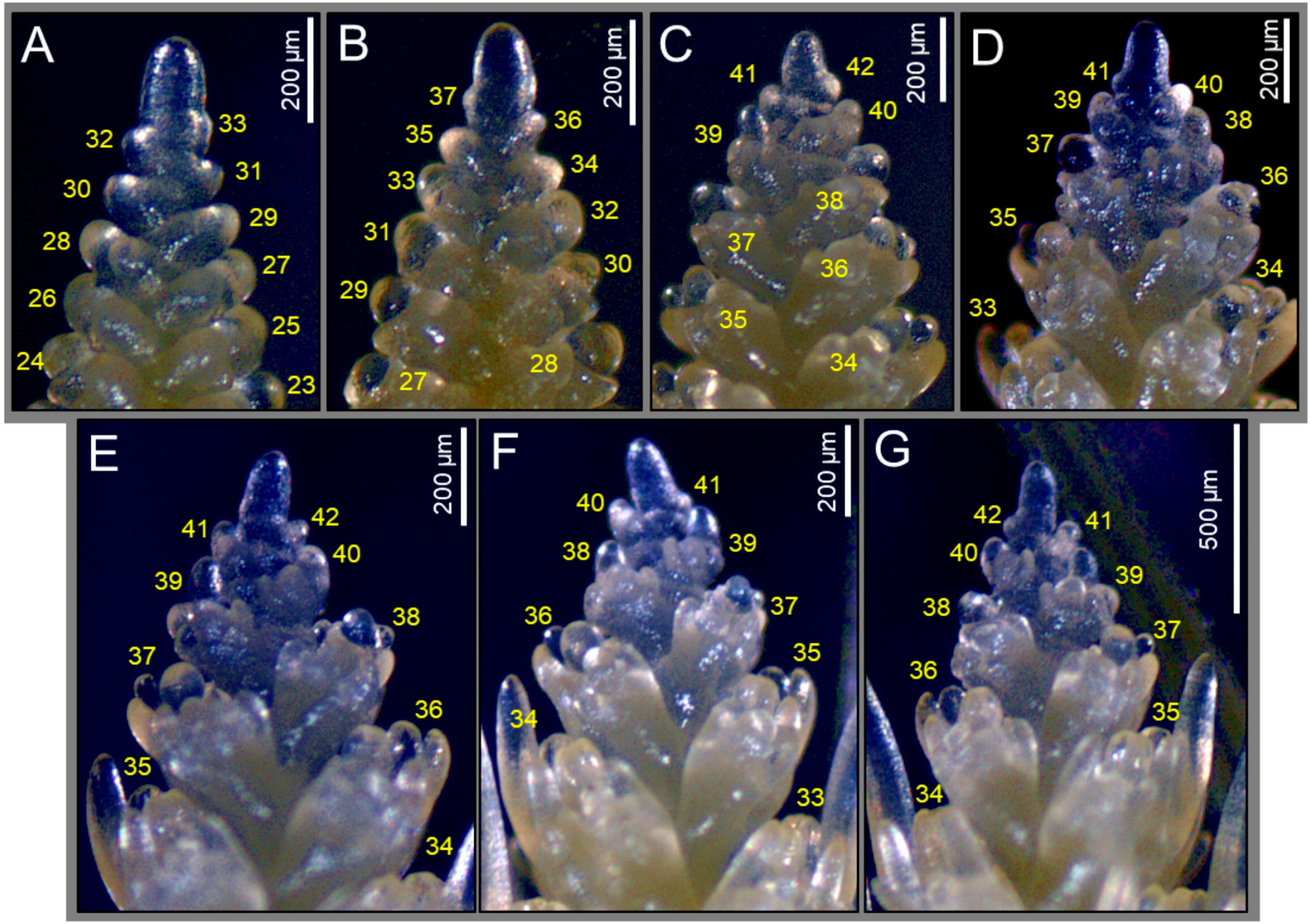
Arrest of spikelet initiation marks the maximum yield potential (MYP) stage in barley spikes. This figure displays the apices of the representative spikes of stages W4.0 to W7.0, shown in Figure 2. The spike apex or the inflorescence meristem (IM) of W4.0 is shown in (A). It is smooth and had a rounded tip. The IM of W4.5 appeared similar to W4.0 (B), while the IM of W5.0 was deformed, and it lost the rounded tip (C). Interestingly, the number of spikelet ridges were proliferated only until W5.0, which coincided with the IM deformation. This suggests that the IM lost its activity before W5.0, resulting in the arrest of spikelet ridges. After W5.0, the IM of consecutive stages (W5.5 to W7.0) (D-G) never regained the rounded tip; instead, their structures were further deformed, and they become translucent. Thus, this palette of images indicated that W5.0 is the MYP stage. W-Waddington scale.

**Figure 4.**
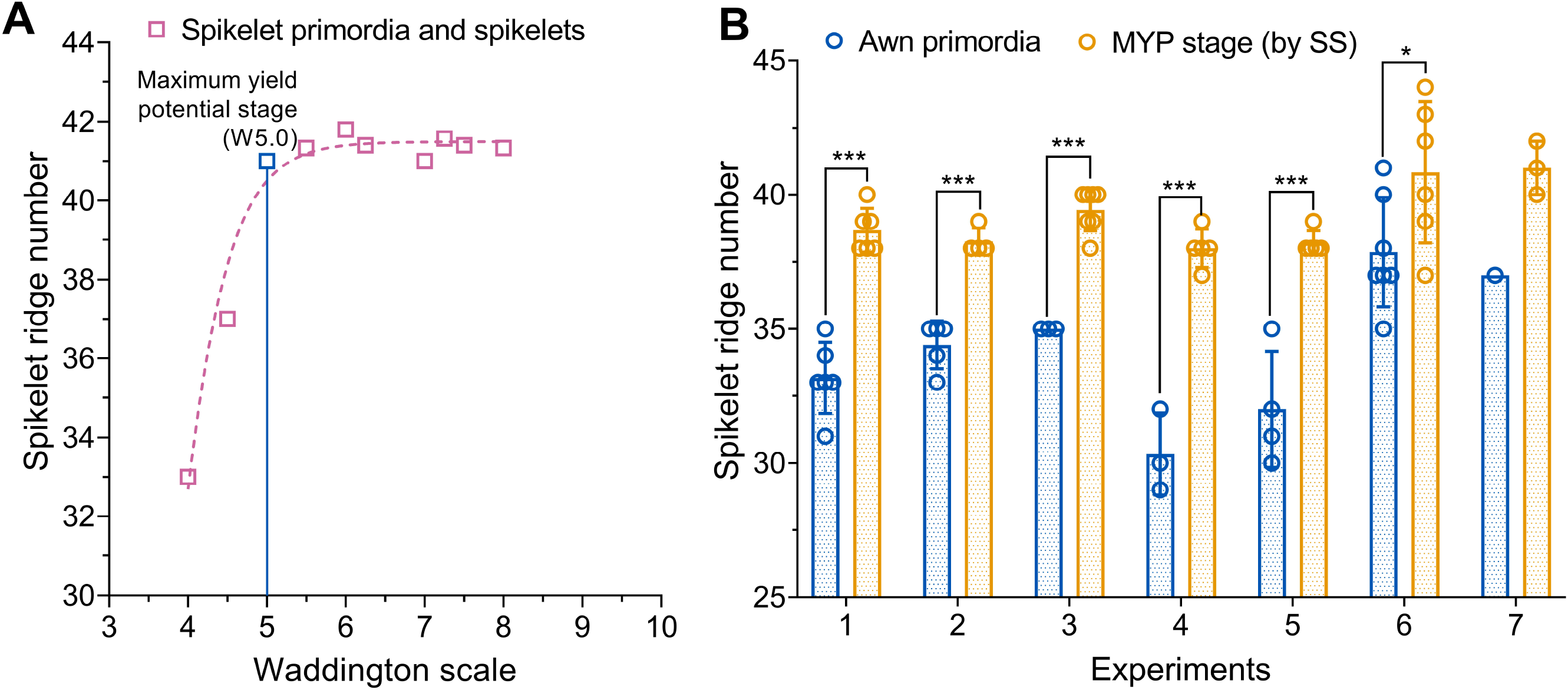
The maximum yield potential (MYP) stage can vary depending on growth conditions. An example of the identification of the MYP stage from experiment 7 is shown in (A). Here, the graph’s inflection point is the MYP stage of this spike. In (B), we showed the comparison of spikelet ridge numbers counted at awn primordium stage (AP, W4.5) and MYP stage (by spikelet stop, SS) from all the seven experiments. The graph points out that in almost all the experiments (except experiment 7), the spikelet ridge number counted at the AP stage is significantly lower than the MYP stage. Due to insufficient replicates of the AP stage in experiment 7, the significance analysis was not performed. Data in A are the mean values of spikelet ridge number counted at various developmental stages; except for the stages W4.0 and W4.5 (AP) (have one replicate), all other stages have three to seven replicates. Data in B were analyzed by multiple Student’s I-tests with false discovery analysis of Benjamini, Kreier, and Yekutieli with the Q value of 5%; except for experiment 7, all other experiments have three to seven replicates; *, P<0.05; **, P<0.001. Replicates are shown as circles and error bars are SD. W-Waddington scale.

### The MYP stage is influenced by genotypic and environmental variation

Based on the results from the Bowman spikelet-tracking experiments, we applied the SS method for the MYP stage identification in two experiments performed in the greenhouse and field with a panel (27 accessions) of two- and six-rowed genotypes (Table S1). In the greenhouse, both row-types had a narrow window for attaining the MYP stage, i.e., as early as ~W4.3 to later stages of ~W6.0 (Fig. 5). In the field, the early MYP stage (~4.5) was not fundamentally different from the greenhouse; however, MYP stages were delayed to ~W7.3 in some accessions (Fig. 5), indicating that soil, light, and temperature affected their development. Six two-rowed (BCC801, BCC929, PI467826, HOR18914, BCC1707, and Bowman) and four six-rowed (BCC881, BCC766, BCC161, and BCC1488) genotypes reached the MYP stage at similar stages (i.e., non-significant difference) in both the growth conditions (Fig. 5A & B). In the field, nine two-rowed (BCC1419, BCC1440, BCC1367, BCC1433, BCC1398, BCC1408, Garnett, BCC1705, and Metcalfe) and three six-rowed (BCC192, BCC719, and Morex) genotypes attained the MYP stage significantly later than the greenhouse (Fig. 5A & B). Interestingly, two two-rowed (HOR2828 and Hockett) and three six-rowed (BCC768, BCC149, and Newdale) genotypes reached the MYP stage significantly earlier in the field than the greenhouse (Fig. 5A & B). A mean value comparison of both row-types’ for MYP stages from the two growth conditions revealed that the two-rowed types reached the MYP stage significantly later in the field than in the greenhouse due to the behavior of seven (BCC1419, BCC1440, BCC1367, BCC1433, BCC1398, Garnett, and Metcalfe) genotypes. They appeared to be as a sub-population within the two-rowed panel (Fig. 5C). Interestingly, among the 27 genotypes grown in two environments, only three six-rowed genotypes (BCC149 and BCC161 in the field; BCC1488 in the greenhouse) attained the MYP stage at or around the AP stage (~W4.5) (Fig. 5B). We also measured the timing to reach the MYP stage in both the experiments by calculating the growing degree days (GDDs). From the analysis of GDDs, it was found that six of the nine two-rowed genotypes, which had a higher MYP in the field, also took significantly longer GDDs to reach the MYP stage in the field (Fig. 6A). In six-rowed, all the three genotypes with a higher MYP in the field availed more GDDs to get to the MYP stage (Fig. 6B). The mean value comparison of GDDs in the two growth conditions showed that only in the field the two-rowed panel got separated into two sub-panels (Fig. 6C) as with the MYP stage (Fig. 5C). From these results, it becomes clear that the MYP stage can be influenced by genotype and growth conditions.

**Figure 5:**
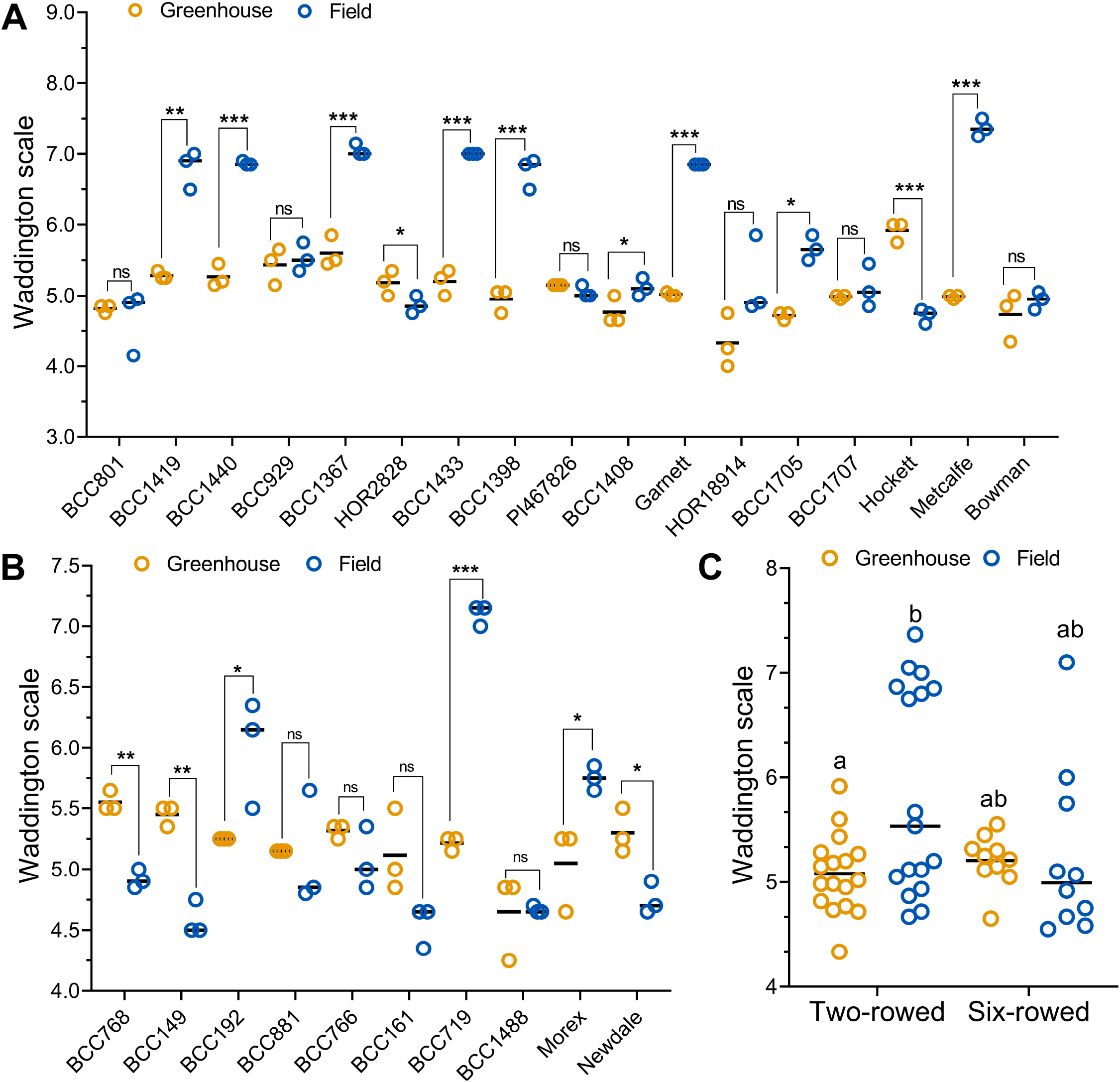
Maximum yield potential (MYP) stage is influenced by genotypic and environmental variation. The MYP stage of 17 two-rowed (A) and ten six-rowed (B) barleys grown in the greenhouse and field plotted according to the Waddington scale. Also, a comparison of the MYP stage mean value of the two- and the six-rowed panel is shown in C. In the greenhouse, 16 out of the 17 two-rowed and all the six-rowed types reached the MYP stage between W4.5 to W5.5. However, in the field, eight genotypes from the two-rowed and three from the six-rowed reached the MYP stage relatively late between W6.0 to W8.0. Interestingly, one of the two-rowed genotypes, HOR2828, and three six-rowed genotypes, reached its MYP stage earlier in the field. Each genotype was represented by three different plants in both the greenhouse and field experiments. W-Waddington scale. Data in A & B were analyzed by multiple Student’s t-tests with false discovery analysis of Benjamini, Kreier, and Yekutieli with the Q value of 5%; *, P<0.05; **, P<0.01; ***, P<0.001; ns, non-significant. Data in C were analyzed by a two-way ANOVA with Tukey’s multiple comparison test (alpha=5%). Different letters denote the statistical difference of adjusted P<0.05.

**Figure 6:**
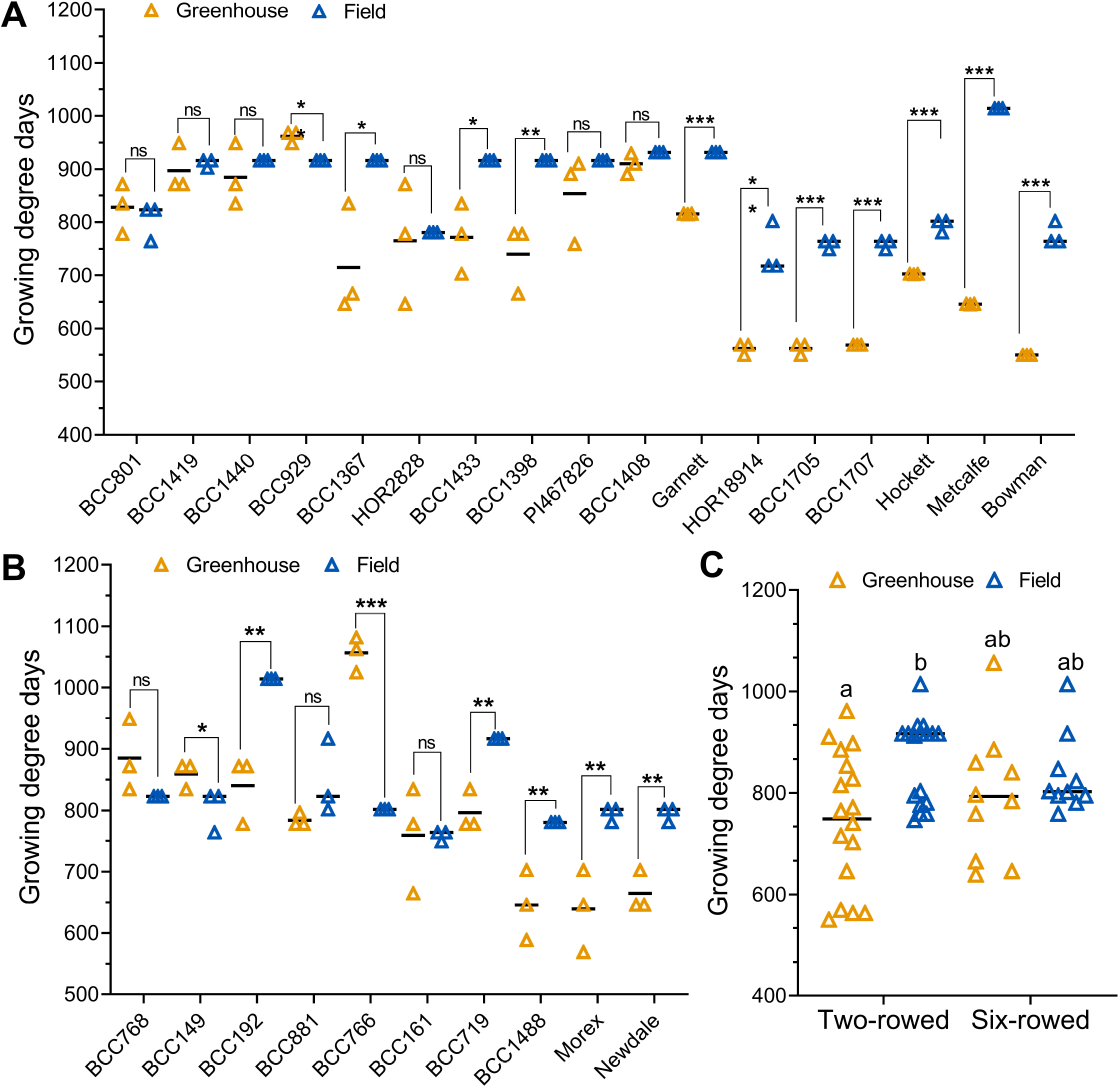
Timing to reach the maximum yield potential (MYP) stage is influenced by genotypic and environmental variation. We show the variation of growing degree days (GDDs) to get to the MYP stage of 17 two-rowed (A) and ten six-rowed (B) barleys grown in the greenhouse and field. A mean value comparison of GDDs to get to the MYP stage of both the panel is shown in C. In the greenhouse, both the panels attained the MYP stage between 550 to 1000 GDDs, while in the field, from 750 to 1050 GDDs. Ten two-rowed and five six-rowed types took more GDDs to reach the MYP stage in the field than the greenhouse. A two-rowed genotype BCC929 (A) and a six-rowed genotype BCC149 (B) made it to the MYP stage earlier in the field than the greenhouse. Each genotype was represented by three different plants in both the greenhouse and field experiments. Data in A & B were analyzed by multiple Student’s t-tests with false discovery analysis of Benjamini, Kreier, and Yekutieli with the Q value of 5%; *, P<0.05; **, P<0.01; ***, P<0.001; ns, non-significant. Data in C were analyzed by a two-way ANOVA with Tukey’s multiple comparison test (alpha=5%). Different letters denote the statistical difference of adjusted P<0.05.

### Timing and stage of the MYP may determine the grain number in two-rowed barley

To better understand the influence of the timing and stage of the MYP in relation to the final main culm grain number (GN), we evaluated the interaction of GDDs to reach the MYP stage and the developmental stage at which the MYP is reached with potential spikelet number (PSN), final spikelet number (SN) and GN of the main culm (Fig.7 & 8). For both row-types, the SRN at the MYP stage was multiplied by three to get the PSN because barley forms three spikelets at every rachis node (Bonnett, 1935; Komatsuda et al., 2007). By combining the results of both the growth conditions, it was clear that the two-rowed PSN, SN, and GN are strongly determined by the time taken to get to the MYP stage (Fig. 7A-C). However, the same traits analyzed for six-rowed types appeared independent of the GDDs (Fig. 7D-F). Also, the MYP stage (as Waddington scale) of the two-rowed significantly influenced the GDDs, PSN, SN, and GN (Fig. 8A-D), while in the six-rowed, only the GDDs were moderately influenced by the MYP stage (Fig. 8E-H). Thus, our interaction study suggests that the timing and developmental stage at which two-rowed barleys reach their MYP stage may determine their main culm yield potential, whereas this is less valid for six-rowed barleys.

**Figure 7:**
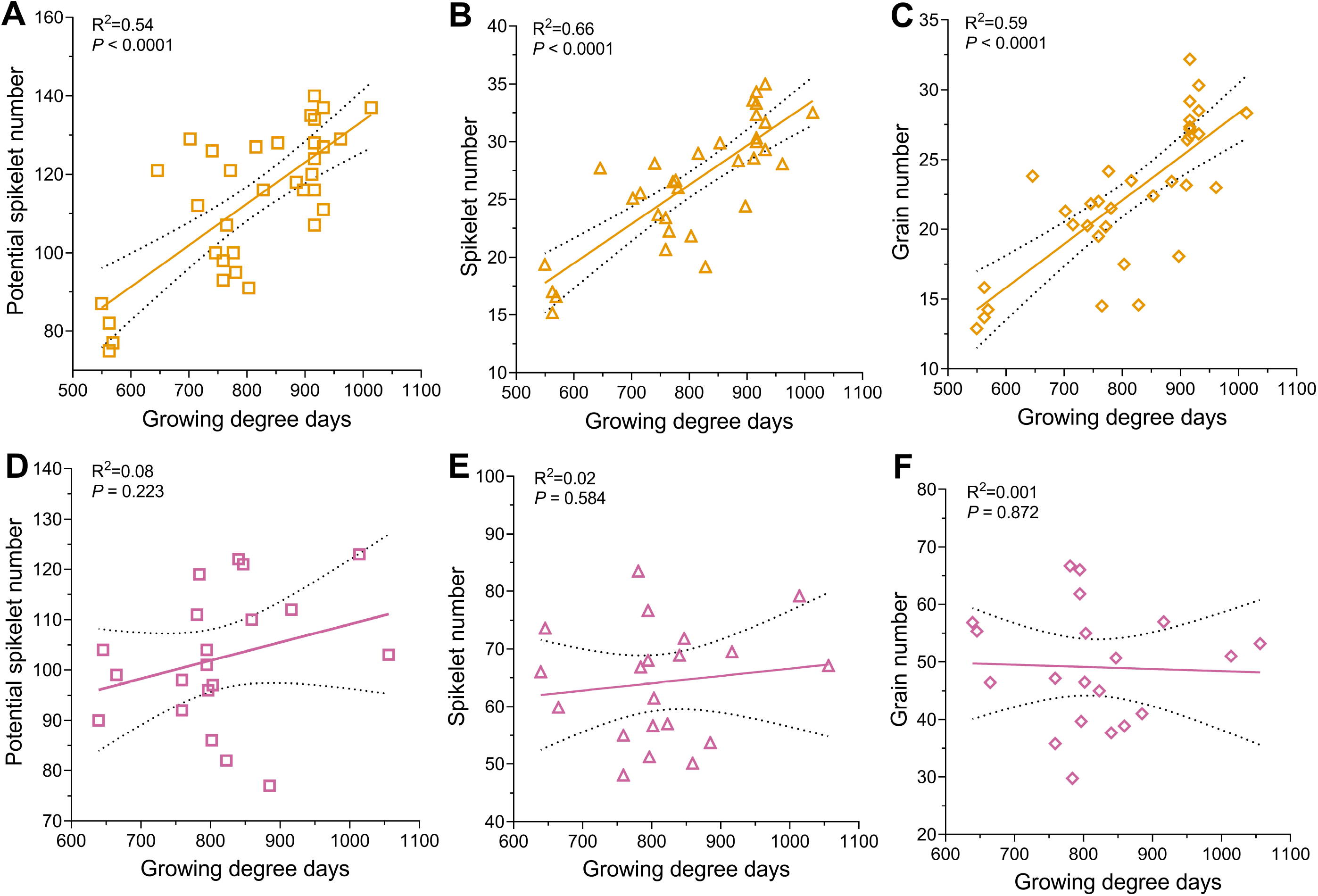
Interactions of timing to the maximum yield potential (MYP) stage and yield components. The growing degree days (GDDs) taken to reach the MYP stage and its interaction with potential spikelet number (PSN), spikelet number (SN), and grain number (GN) of two-(A, B, C) and six-rowed types (D, E, F) grown in the greenhouse and field are shown. In two-rowed, genotypes that spent more GDDs to get to the MYP stage developed more PSN (A), SN (B), and GN (C), while six-rowed genotypes did not depend on the GDDs to MYP stage for the production of PSN (D), SN (E), and GN (F). GDDs and PSN were taken from three different plants, while SN and GN were from six other plants. The 95% confidence intervals were identified for every linear regression and plotted as confidence bands (black dotted line) along with the ‘goodness of fit’ line.

**Figure 8:**
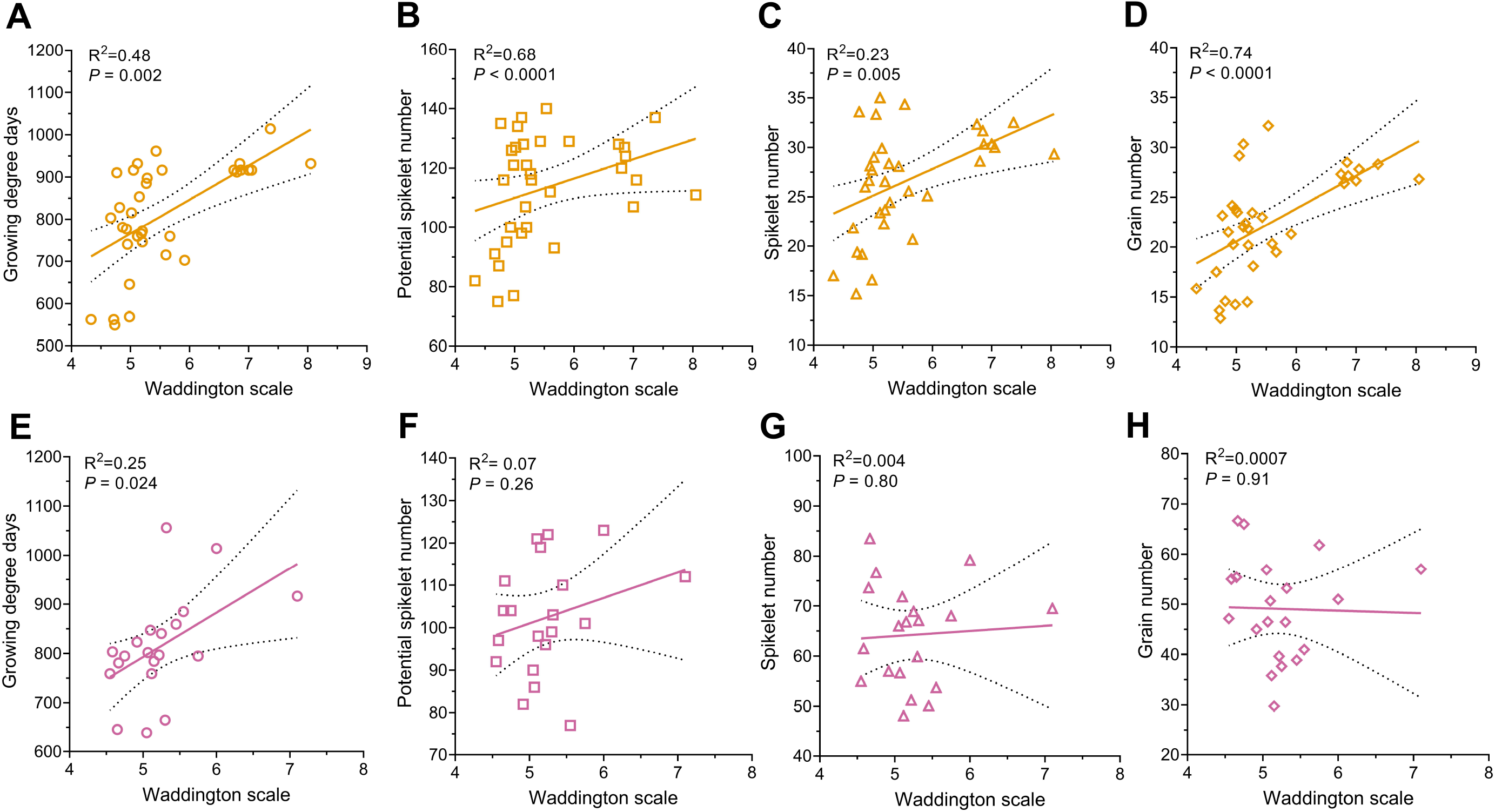
Interactions of the maximum yield potential (MYP) stage and yield components. The MYP stage (as Waddington scale) and its interaction with growing degree days (GDDs) required to reach the MYP stage, potential spikelet number (PSN), spikelet number (SN), and grain number (GN) of two-(A, B, C, D) and six-rowed types (E, F, G, H) grown in the greenhouse and field are shown. In both row-type panels (greenhouse and field), genotypes reached the MYP stage at a later stage of the Waddington scale, taking more GDDs than earlier genotypes (A&E). In two-rowed, genotypes get to MYP at a later stage of the Waddington scale produced more PSN (B), SN (C), and GN (D), while six-rowed types’ PSN (E), SN (F), and GN (G) are independent of the MYP stage. GDDs and PSN were taken from three different plants, while SN and GN were from six other plants. The 95% confidence intervals were identified for every linear regression and plotted as confidence bands (black dotted line) along with the ‘goodness of fit’ line.

## Discussion

Improvement of floret/spikelet fertility can be a promising avenue to enhance the yield of small grain cereals. In general, the floret/spikelet fertility or efficiency of floret/spikelet survival is measured by comparing the maximum yield potential (MYP) with the final floret/spikelet number. Developing a method to identify the barley MYP would pave the way to understand the genetics and mechanism of spikelet survival in barley. From the spikelet initiation and growth study with the two-rowed cv. Bowman and comparing the MYP stages in a panel of 27 barley accessions, we found that the MYP stage can be different from the AP stage and that its observed variation mainly depends on genetic and environmental attributes. Also, we propose that the shape and activity of the IM can be used as a proxy to identify the end of the spikelet initiation or the MYP stage. We also discovered that the GDDs to reach the MYP stage, as well as the MYP stage of two-rowed genotypes, may determine the final spikelet and grain number.

### The arrest of spikelet initiation specifies the MYP stage in barley, and it can be different from the AP stage

The concept behind the identification of the MYP stage in barley is that it helps to assess how efficiently a plant produces grains from its maximum number of spikelets. To this end, it is crucial to recognize the stage at which a spike possesses its maximum number of spikelets. Several reports evaluated the rate of barley spikelet development and identified the MYP stage by following the formation of spikelets until the cessation of spikelet initiation (Gallagher et al., 1976; Kirby, 1977; Arisnabarreta and Miralles, 2006a; Arduini et al., 2010). However, reports aimed to reveal the genetic variation of barley floret/spikelet fertility (Alqudah and Schnurbusch, 2014) or responses to different treatments and growth conditions (Arisnabarreta and Miralles, 2004, 2006a; Arisnabarreta and Miralles, 2010, 2015) assumed the AP stage as the MYP in all the genotypes and different environmental conditions. Our comprehensive study using the two-rowed cv. Bowman revealed different spikelet initiation and growth phases, as previously shown for cereals (Gallagher, 1979; Appleyard et al., 1982). The end of spikelet number increase, or the first inflection point of the graph (Fig.1
), denoted the cessation of the spikelet initiation phase at which a spike possesses its maximum number of spikelets, known as the MYP stage. Interestingly, in all our experiments with the cultivar Bowman, the MYP stage of the spike was significantly different from the AP stage (~W4.5) (Fig. S2 A & B) contrasting to the previous report (Appleyard et al., 1982). Similarly, other studies reported that the culmination of spikelet initiation occurred concurrently with stages of stamen primordia (W3.5) (Nicholls, 1963; Nicholls, 1974), W6.0 (shortly after the initiation of the pistil) (Waddington et al., 1983) and after AP (Arduini et al., 2010). In wheat, a study conducted with 12 genotypes reported that the MYP stage consistently occurred during the green anther (GA) stage (Kirby and Appleyard, 1984) among the genotypes; but similarly in the tiller removal treatment (Guo and Schnurbusch, 2015). However, a closer look at the data indicated that in three instances of detillered plants, the MYP stage and the number of florets were significantly different from the GA stage (Guo & Schnurbusch, 2015; Table 3), somewhat underestimating the maximum floret number most likely due to a treatment effect (Fig. S3). The above studies suggest that, regardless if in wheat or barley, researchers should be cautious when analyzing the MYP stage in a limited number of genotypes because most likely, the difference in the MYP stage is subtle and/or found only in a few genotypes. Our MYP stage comparison study with the panel (27 genotypes) of two- and six-rowed barley types clearly shows that the MYP stage can be attained from ~W4.0 to ~W7.5 (Fig. 5) by different genotypes in variable growth conditions. Furthermore, tracking the spike apex during early spike development unveiled that the IM lost its meristematic activity, visualized by its deformed dome at W5.0 (Fig. 3C) as reported previously (Kirby, 1977). All these findings demonstrate that, in barley, spikelet initiation arrest marks the MYP stage, which can be different from the AP stage.

### MYP stage is plastic and may influence the GN of two-rowed barleys

Our extensive studies on Bowman spikelet initiation and growth and the comparison of MYP stages on a selected panel exemplified that the MYP stage is influenced by the genetic variation and different growth conditions. Furthermore, these studies also indicated that spikelet ridge number (SRN) were significantly different in the AP and MYP stage (identified by SS) (Fig. 4B & S4). Intriguingly, the mean SRN counted at the MYP stage (identified by SS) of all the seven Bowman experiments showed a narrow range, 38 to 41; however, a few experiments have similar MYP, and some are significantly different from each other (Fig. S2C). Similarly, the MYP comparison study on the selected panel showed 12 out of 27 genotypes had similar SRN in both the growth conditions (Fig. S5). Interestingly, four among the above 12 genotypes and six more genotypes (in total, ten) had a similar spike developmental scale when they were at the MYP stage (Fig. 5A & B). The above results also implied that 15 and 17 (out of 27) genotypes had significantly different SRN and developmental scales, respectively, when they got to the MYP stage (Fig. S4, 5A & B). A few studies using a limited number of barley genotypes (from four to ten) claimed that the maximum number of spikelet primordia was not affected due to lower nitrogen supply (Arisnabarreta and Miralles, 2004), shading (Arisnabarreta and Miralles, 2008b), and different growth seasons (Arisnabarreta and Miralles, 2006a). However, there are also studies with a few accessions (three to six) that reported that the MYP varies among different accessions and sowing dates (Arduini et al., 2010; Digel et al., 2015). Furthermore, our GDDs analysis of the MYP stage on the selected panel revealed that analogous to the MYP and its developmental stage, there were two groups: one group (nine genotypes) that did not show a significant difference in GDDs between the growth conditions; whereas other genotypes (18) had significant differences (Fig 6). Based on our findings and the above reports, we propose that the MYP stage, maximum SRN, and GDDs to MYP stage are primarily dependent on the genotypic variation and environmental conditions. Our regression analysis of GDDs to MYP and MYP stage with the yield potential traits on the selected panel suggests that timing to reach the stage of MYP in two-rowed strongly influences GN of the main culm (Fig. 7 & 8). A similar finding was reported for two-rowed genotypes that had different alleles of *Ppd-H1*, where, under long-day conditions, introgression lines with a mutant *ppd-H1* took a longer duration to reach W3.5 and produced more grains per spike compared to the barley lines with the wildtype allele (Digel et al., 2015). Nevertheless, we propose that this association of MYP and GN of two-rowed genotypes must be verified with a bigger panel of accessions representing the available variations in barley. Thus, our study’s findings suggest that the MYP stage and its associated traits are plastic, and it may influence the yield potential of two-rowed barley accessions.

## Conclusion

Identifying the MYP stage and finding the maximum number of spikelets of a barley spike are two different tasks. Determining the MYP stage requires tracking the spikelet initiation from the early development; therefore, it is almost impossible to do that for a bigger panel (several hundred) of barley accessions. From the knowledge gained from this study, as well as from the previous publications, we, therefore, recommend a few key steps to identify the MYP of a barley spike easily. An initial spike dissection (shown in Kirby et al., 1984) must be performed when the first node (Zadoks et al., 1974) is detected on the main culm to know the spike developmental stage. A study also reported that a few barley genotypes reached their MYP around this stage (Arduini et al., 2010). However, for a bigger panel of accessions, one would not get the MYP for all genotypes at the first node stage. Thus, it may be necessary to go for another one or two times dissection. We suggest that the second dissection could be after a week or two depending on the rate of development. Suppose, at the second dissection, if the spike already entered into the spikelet degeneration (abortion) phase, it would be still possible to count the spikelet ridges, because during early stages of abortion the spike apex does not disintegrate (Fig. S6). To verify whether the spike already reached the MYP, the change of the IM shape (tapered IM) (Fig.3) can be used as a visible marker. If the spike at the second dissection is not in the MYP stage, a third attempt similar to the second dissection could be tried. Based on our experience from this study, we believe that an early first dissection (around the first node detection) followed by a periodic one or two dissections might help to get the MYP of most of the genotypes. Notably, the plateau stage of spike development may provide an excellent opportunity to assess the MYP of variable genotypes at varying stages.

## Supplementary data

Table S1: Details of the 27 accessions used in this study.

Table S2: Growth conditions of the field and greenhouse experiments.

Table S3: Growth conditions of the Bowman experiments.

## Acknowledgments

We are sincerely thankful to Angelika Püschel, Corinna Trautewig, and Mechthild Pürschel for their excellent technical support; Ravi Koppolu, Nandhakumar Shanmugaraj, and Roop Kamal for their help with sampling and harvesting. The authors are grateful to Ravi Koppolu, Roop Kamal, and Yongyu Huang for critically reading a previous version of the manuscript. For conducting this study T.S. received financial support from the European Research Council (ERC), grant agreement 681686 “LUSH SPIKE,” ERC-2015-CoG; the HEISENBERG Program of the German Research Foundation (DFG), grant nos. SCHN 768/8-1 and SCHN 768/15-1 and the IPK core budget.

## Author contributions

T.S. conceived the project and supervised the study. T.S. and V.T. conceived and designed the idea for this study, while V.T. developed the ‘Spikelet Stop’ method for determining the MYP stage; V.T. performed the experiments, analyzed the data, and wrote the manuscript; T.S. revised and reviewed the manuscript.

## Supplementary figure legends

Figure S1: **Carpel development from W4.5 to W7.0.** Development of carpels according to the scale proposed by Waddington et al., 1983 is given in A-F. Figure A indicates the Waddington (W) stage 4.5, B shows W5.0, C displays W5.5, D specifies W6.0, E denotes W6.25, and F discloses W7.0. W-Waddington scale.

Figure S2: **Maximum yield potential (MYP) stage can be different from the awn primordium stage.** The Waddington scales of the maximum yield potential (MYP) stage (by spikelet stop, SS) and the awn primordium stage (AP) identified in seven experiments are shown in (A), and the Waddington scales of the MYP stage (by SS) found in the seven experiments are shown in (B). The variation of spikelet ridge number at the MYP stage (by SS) in all the seven experiments is shown in (C). Except for experiment 7 that had only one replication for the AP stage, all other experiments had three to seven replications. Data in A were analyzed by multiple *Student’s t*-tests with false discovery analysis of *Benjamini, Kreier*, and Yekutieli with the Q value of 5%; ***, *P*<0.001. Data in C were analyzed by a one-way ANOVA with Tukey’s multiple comparison test (alpha=5%). Different letters denote the statistical difference of adjusted *P*<0.05. Replicates are shown as circles and error bars are SD.

Figure S3: **Reanalysis of the de-tillered data from Guo & Schnurbusch, 2015.** By reanalyzing the table 3 data (de-tillering experiment), we found that in three instances, the maximum yield potential (MYP) stage is significantly different from the GA (green anther) stage. Importantly, this data represents a single spikelet located in various positions (central, basal, or apical). If we extrapolate this discrepancy to the whole spike, it indicates that one might underestimate the MYP of certain genotypes. Data are shown as mean±SD; n=3; Data was analyzed by the *Student’s two-tailed t-test; ***,* P<0.001; **, P<0.01; *, P<0.05.

Figure S4: **Maximum yield potential (MYP) stage can be different from the awn primordium stage.** We show the number of spikelet ridges counted for various genotypes at the awn primordium (AP) stage and maximum yield potential (MYP) stage (by spikelet stop, SS) in the greenhouse (A) and field (B). From the 13 genotypes’ available data, only one two-rowed type BCC1367 had a similar number of spikelet ridges both in the AP and MYP stage (by SS). Each genotype was represented by three different plants in both the greenhouse and field experiments. Data in A & B were analyzed by multiple *Student’s t*-tests with false discovery analysis of *Benjamini, Kreier*, and Yekutieli with the Q value of 5%; *, *P*<0.05; **, *P*<0.01; ***, *P*<0.001. Replicates are shown as circles and error bars are SD.

Figure S5: **Maximum yield potential comparison of 27 barley accessions.** We show the variation of maximum yield potential (MYP) as spikelet ridge number (SRN) in 17 two-rowed (A) and ten six-rowed (B) barleys grown in the greenhouse and field. A mean value comparison of the MYP of both the panel is shown in C. Each genotype was represented by three different plants in both the greenhouse and field experiments. Data in A & B were analyzed by multiple *Student’s t*-tests with false discovery analysis of *Benjamini, Kreier*, and Yekutieli with the Q value of 5%; *, *P*<0.05; **, *P*<0.01; ***, *P*<0.001; ns, non-significant. Data in C were analyzed by a two-way ANOVA with Tukey’s multiple comparison test (alpha=5%); ns, non-significant.

Figure S6: **Spikelet ridges on aborted spike apices.** We displayed examples of aborted spike apices and their counted spikelet ridges in A, B, & C.

## References

Alqudah, A.M., and Schnurbusch, T. (2014). Awn primordium to tipping is the most decisive developmental phase for spikelet survival in barley. Functional Plant Biology 41, 424.

Alqudah, A.M., Sharma, R., Pasam, R.K., Graner, A., Kilian, B., and Schnurbusch, T. (2014). Genetic dissection of photoperiod response based on GWAS of pre-anthesis phase duration in spring barley. PLoS One 9, e113120.

Appleyard, M., Kirby, E., and Fellowes, G. (1982). Relationships between the duration of phases in the pre-anthesis life cycle of spring barley. Aust J Agr Res 33, 917–925.

Arduini, I., Ercoli, L., Mariotti, M., and Masoni, A. (2010). Coordination between plant and apex development in Hordeum vulgare ssp. distichum. C R Biol 333, 454–460.

Arisnabarreta, S., and Miralles, D.J. (2004). The influence of fertiliser nitrogen application on development and number of reproductive primordia in field-grown two- and six-rowed barleys. Aust J Agr Res 55, 357–366.

Arisnabarreta, S., and Miralles, D.J. (2006a). Floret development and grain setting in near isogenic two- and six-rowed barley lines (Hordeum vulgare L.). Field Crops Research 96, 466–476.

Arisnabarreta, S., and Miralles, D.J. (2006b). Yield Responsiveness in Two- and Six-Rowed Barley Grown in Contrasting Nitrogen Environments. Journal of Agronomy and Crop Science 192, 178–185.

Arisnabarreta, S., and Miralles, D.J. (2008a). Critical period for grain number establishment of near isogenic lines of two-and six-rowed barley. Field Crops Research 107, 196–202.

Arisnabarreta, S., and Miralles, D.J. (2008b). Radiation effects on potential number of grains per spike and biomass partitioning in two- and six-rowed near isogenic barley lines. Field Crops Research 107, 203–210.

Arisnabarreta, S., and Miralles, D.J. (2010). Nitrogen and radiation effects during the active spike-growth phase on floret development and biomass partitioning in 2- and 6-rowed barley isolines. Crop and Pasture Science 61, 578–587.

Arisnabarreta, S., and Miralles, D.J. (2015). Grain number determination under contrasting radiation and nitrogen conditions in 2-row and 6-row barleys. Crop and Pasture Science 66, 456–465.

Bancal, P. (2008). Positive contribution of stem growth to grain number per spike in wheat. Field Crops Research 105, 27–39.

Bancal, P. (2009). Early development and enlargement of wheat floret primordia suggest a role of partitioning within spike to grain set. Field Crops Research 110, 44–53.

Benjamini, Y., Krieger, A.M., and Yekutieli, D. (2006). Adaptive linear step-up procedures that control the false discovery rate. Biometrika 93, 491–507.

Bommert, P., and Whipple, C. (2018). Grass inflorescence architecture and meristem determinacy. Seminars in Cell & Developmental Biology 79, 37–47.

Bonnett, O. (1936). The development of the wheat spike. J agric Res 53, 445–451.

Bonnett, O.T. (1935). The development of the barley spike. J agric Res 51, 451–457.

Bonnett, O.T. (1966). Inflorescences of maize, wheat, rye, barley, and oats: their initiation and development/721. Bulletin (University of Illinois (Urbana-Champaign campus) Agricultural Experiment Station); no 721.

Cottrell, J.E., Easton, R.H., Dale, J.E., Wadsworth, A.C., Adam, J.S., Child, R.D., and Hoad, G.V. (1985). A Comparison of Spike and Spikelet Survival in Mainstem and Tillers of Barley. Annals of Applied Biology 106, 365–377.

del Moral, M.G., Tejada, M.J., Del Moral, L.G., Ramos, J., De Togores, F.R., and Molina-Cano, J. (1991). Apex and ear development in relation to the number of grains on the main-stem ears in spring barley (Hordeum distichon). The Journal of Agricultural Science 117, 39–45.

Digel, B., Pankin, A., and von Korff, M. (2015). Global Transcriptome Profiling of Developing Leaf and Shoot Apices Reveals Distinct Genetic and Environmental Control of Floral Transition and Inflorescence Development in Barley. Plant Cell 27, 2318–2334.

Elhani, S., Martos, V., Rharrabti, Y., Royo, C., and García Del Moral, L.F. (2007). Contribution of main stem and tillers to durum wheat (Triticum turgidum L. var. durum) grain yield and its components grown in Mediterranean environments. Field Crops Research 103, 25–35.

Ferrante, A., Savin, R., and Slafer, G.A. (2012). Floret development and grain setting differences between modern durum wheats under contrasting nitrogen availability. Journal of experimental botany, ers320.

Ferrante, A., Savin, R., and Slafer, G.A. (2013). Is floret primordia death triggered by floret development in durum wheat? J Exp Bot 64, 2859–2869.

Ferrante, A., Savin, R., and Slafer, G.A. (2020). Floret development and spike fertility in wheat: Differences between cultivars of contrasting yield potential and their sensitivity to photoperiod and soil N. Field Crops Research 256, 107908.

Gallagher, J. (1979). Ear development: Processes and prospects. Crop Physiology and Cereal Breeding, 3–9.

Gallagher, J., Biscoe, P., and Scott, R. (1976). Barley and its environment: VI. Growth and development in relation to yield. J Appl Ecol, 563–583.

García, G.A., Dreccer, M.F., Miralles, D.J., and Serrago, R.A. (2015). High night temperatures during grain number determination reduce wheat and barley grain yield: a field study. Global Change Biol 21, 4153–4164.

Gauley, A., and Boden, S.A. (2019). Genetic pathways controlling inflorescence architecture and development in wheat and barley. Journal of Integrative Plant Biology 61, 296–309.

Gonzalez, F.G., Miralles, D.J., and Slafer, G.A. (2011). Wheat floret survival as related to pre-anthesis spike growth. J Exp Bot 62, 4889–4901.

Guo, Z., and Schnurbusch, T. (2015). Variation of floret fertility in hexaploid wheat revealed by tiller removal. Journal of experimental botany 66, 5945–5958.

Kirby, E.J.M. (1977). The Growth of the Shoot Apex and the Apical Dome of Barley During Ear Initiation. Ann Bot-London 41, 1297–1308.

Kirby, E.M., and Appleyard, M. (1984). Cereal development guide. Cereal development guide 2nd Edition.

Komatsuda, T., Pourkheirandish, M., He, C., Azhaguvel, P., Kanamori, H., Perovic, D., Stein, N., Graner, A., Wicker, T., Tagiri, A., et al. (2007). Six-rowed barley originated from a mutation in a homeodomain-leucine zipper I-class homeobox gene. Proc Natl Acad Sci U S A 104, 1424–1429.

Koppolu, R., Anwar, N., Sakuma, S., Tagiri, A., Lundqvist, U., Pourkheirandish, M., Rutten, T., Seiler, C., Himmelbach, A., Ariyadasa, R., et al. (2013). Six-rowed spike4 (Vrs4) controls spikelet determinacy and row-type in barley. Proc Natl Acad Sci U S A 110, 13198–13203.

Koppolu, R., and Schnurbusch, T. (2019). Developmental pathways for shaping spike inflorescence architecture in barley and wheat. Journal of integrative plant biology.

McKim, S.M., Koppolu, R., and Schnurbusch, T. (2018). Barley inflorescence architecture. In The Barley Genome (Springer), pp. 171–208.

McMaster, G.S., and Wilhelm, W. (1997). Growing degree-days: one equation, two interpretations. Agricultural and Forest Meterology 87, 291–300.

Miralles, D.J., Richards, R.A., and Slafer, G.A. (2000). Duration of the stem elongation period influences the number of fertile florets in wheat and barley. Functional Plant Biology 27, 931–940.

Motulsky, H.J., and Brown, R.E. (2006). Detecting outliers when fitting data with nonlinear regression–a new method based on robust nonlinear regression and the false discovery rate. BMC bioinformatics 7, 123.

Nicholls, P. (1963). Studies on the Growth of the Barley Apex I. Interrelationships Between Primordium Formation, Apex Length,. and Spikelet Development. Aust J Biol Sci 16, 561–571.

Nicholls, P.B. (1974). Interrelationship between Meristematic Regions of Developing Inflorescences of Four Cereal Species. Ann Bot-London 38, 827–837.

Peltonen-Sainio, P., Kangas, A., Salo, Y., and Jauhiainen, L. (2007). Grain number dominates grain weight in temperate cereal yield determination: evidence based on 30 years of multi-location trials. Field Crops Research 100, 179–188.

Prystupa, P., Savin, R., and Slafer, G.A. (2004). Grain number and its relationship with dry matter, N and P in the spikes at heading in response to N× P fertilization in barley. Field Crops Research 90, 245–254.

Sakuma, S., Golan, G., Guo, Z., Ogawa, T., Tagiri, A., Sugimoto, K., Bernhardt, N., Brassac, J., Mascher, M., Hensel, G., et al. (2019). Unleashing floret fertility in wheat through the mutation of a homeobox gene. Proceedings of the National Academy of Sciences 116, 5182–5187.

Sakuma, S., and Schnurbusch, T. (2020). Of floral fortune: tinkering with the grain yield potential of cereal crops. New Phytologist 225, 1873–1882.

Serrago, R.A., Alzueta, I., Savin, R., and Slafer, G.A. (2013). Understanding grain yield responses to source–sink ratios during grain filling in wheat and barley under contrasting environments. Field Crops Research 150, 42–51.

Ugarte, C., Calderini, D.F., and Slafer, G.A. (2007). Grain weight and grain number responsiveness to pre-anthesis temperature in wheat, barley and triticale. Field Crops Research 100, 240–248.

Waddington, S.R., Cartwright, P.M., and Wall, P.C. (1983). A Quantitative Scale of Spike Initial and Pistil Development in Barley and Wheat. Ann Bot-London 51, 119–130.

Whipple, C.J. (2017). Grass inflorescence architecture and evolution: the origin of novel signaling centers. New Phytologist 216, 367–372.

Zadoks, J.C., Chang, T.T., and Konzak, C.F. (1974). A decimal code for the growth stages of cereals. Weed Res 14, 415–421.

