## Supplementary figures and images for "A development guide for evaluating the maximum yield potential stage in barley"

### Figure-S1

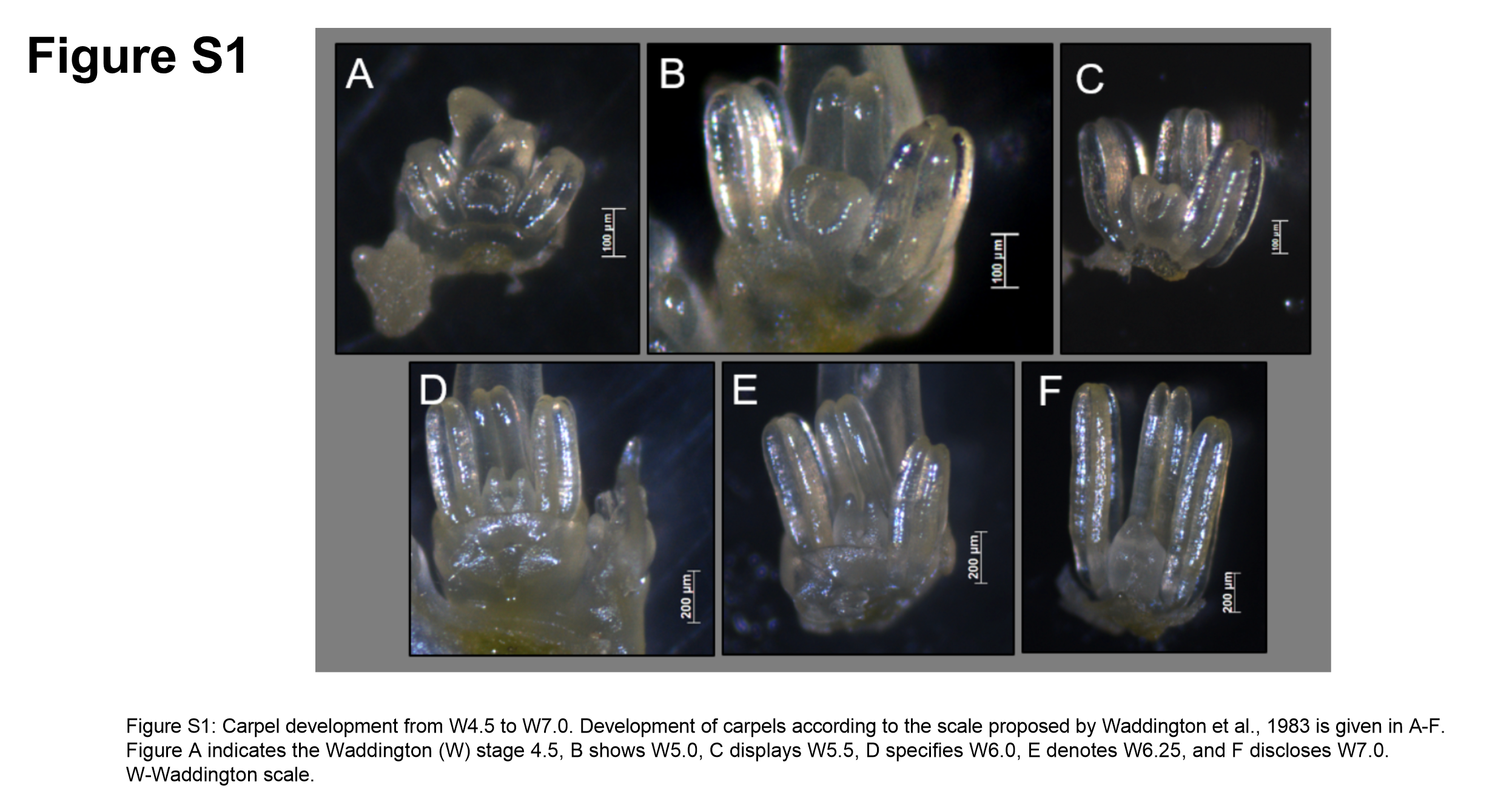

### Figure-S2

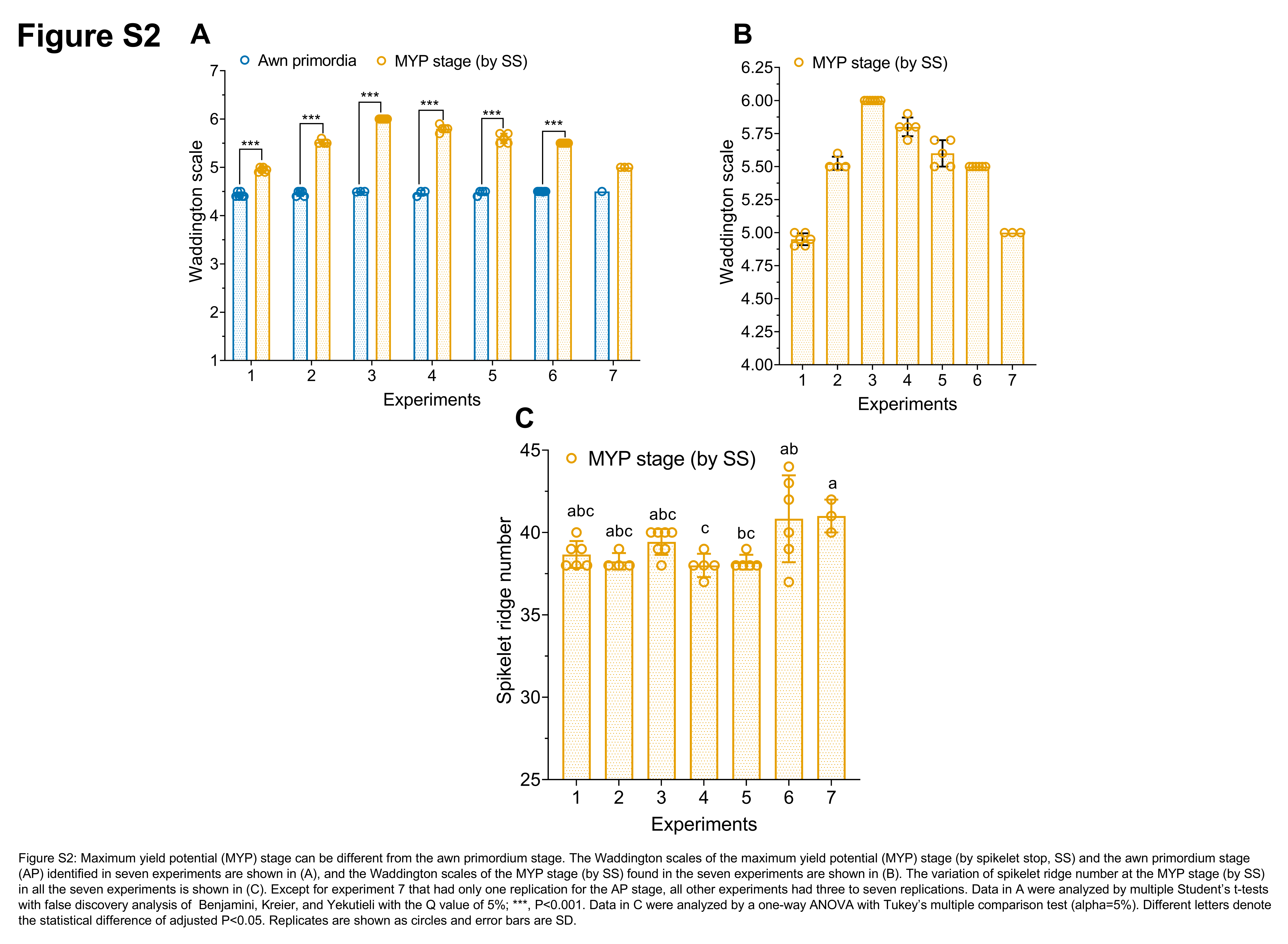

### Figure-S3

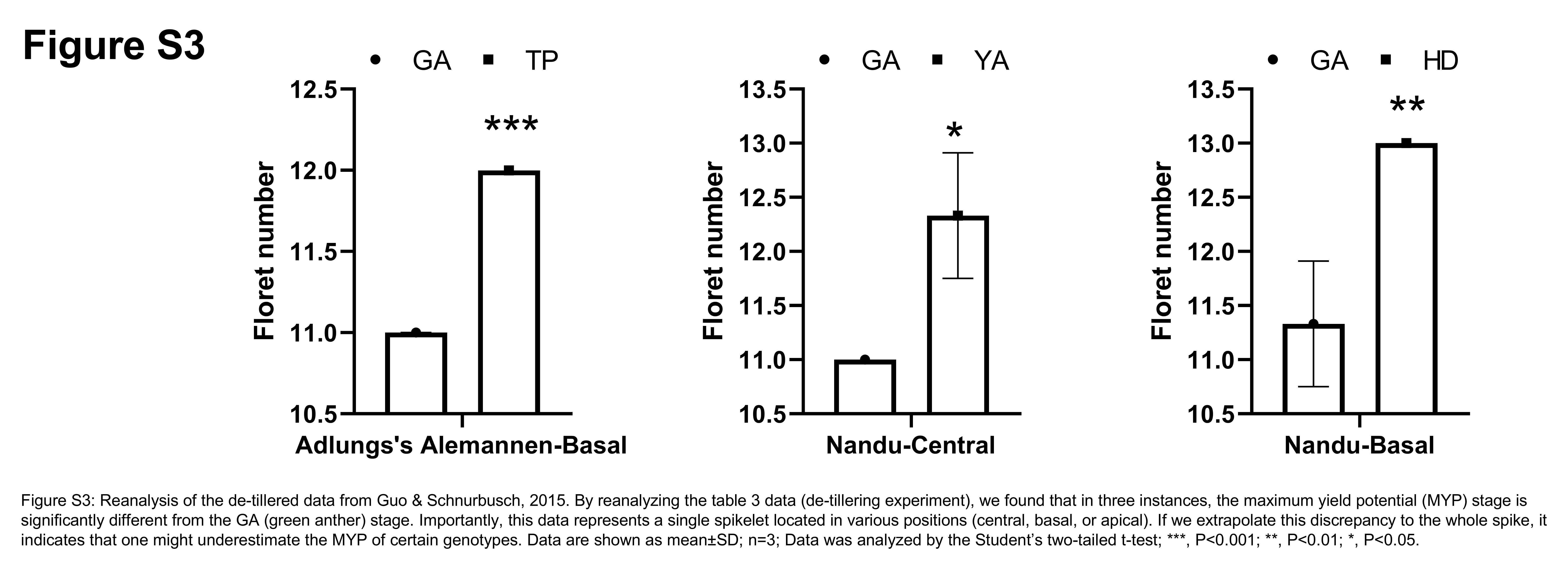

### Figure-S4

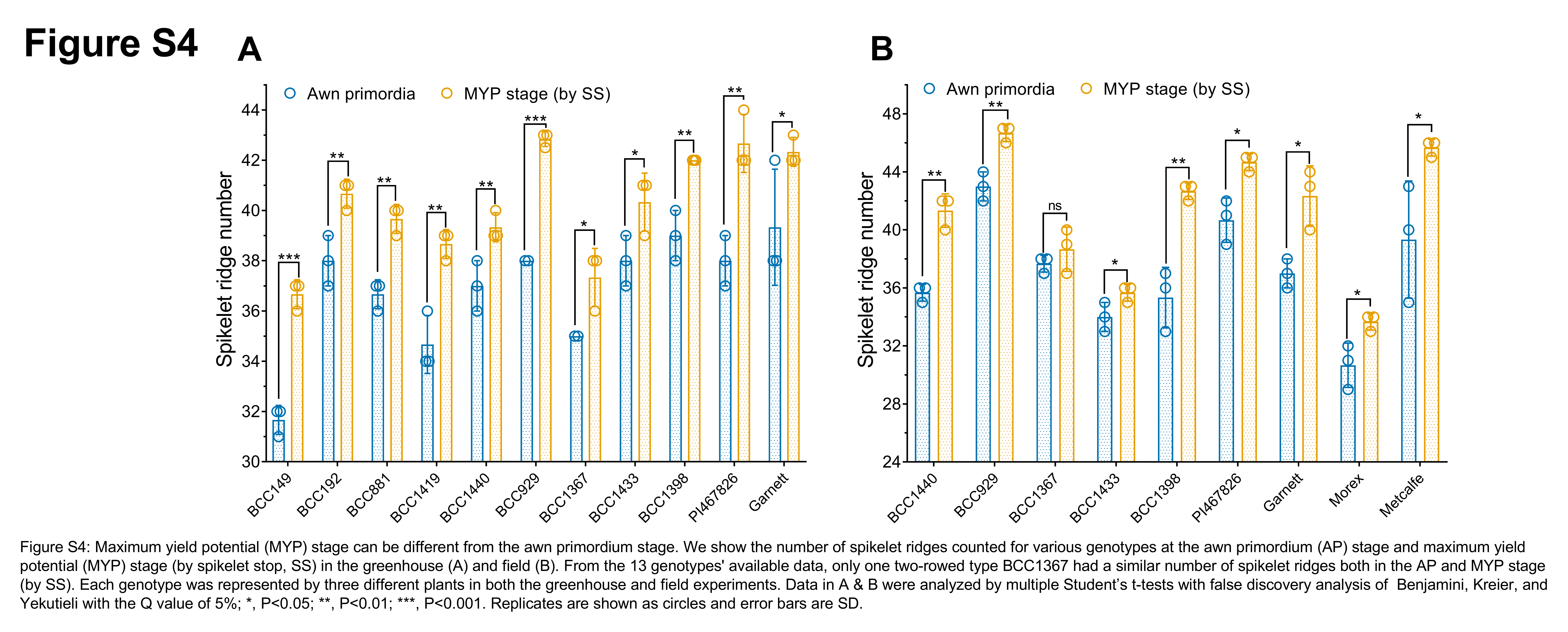

### Figure-S5

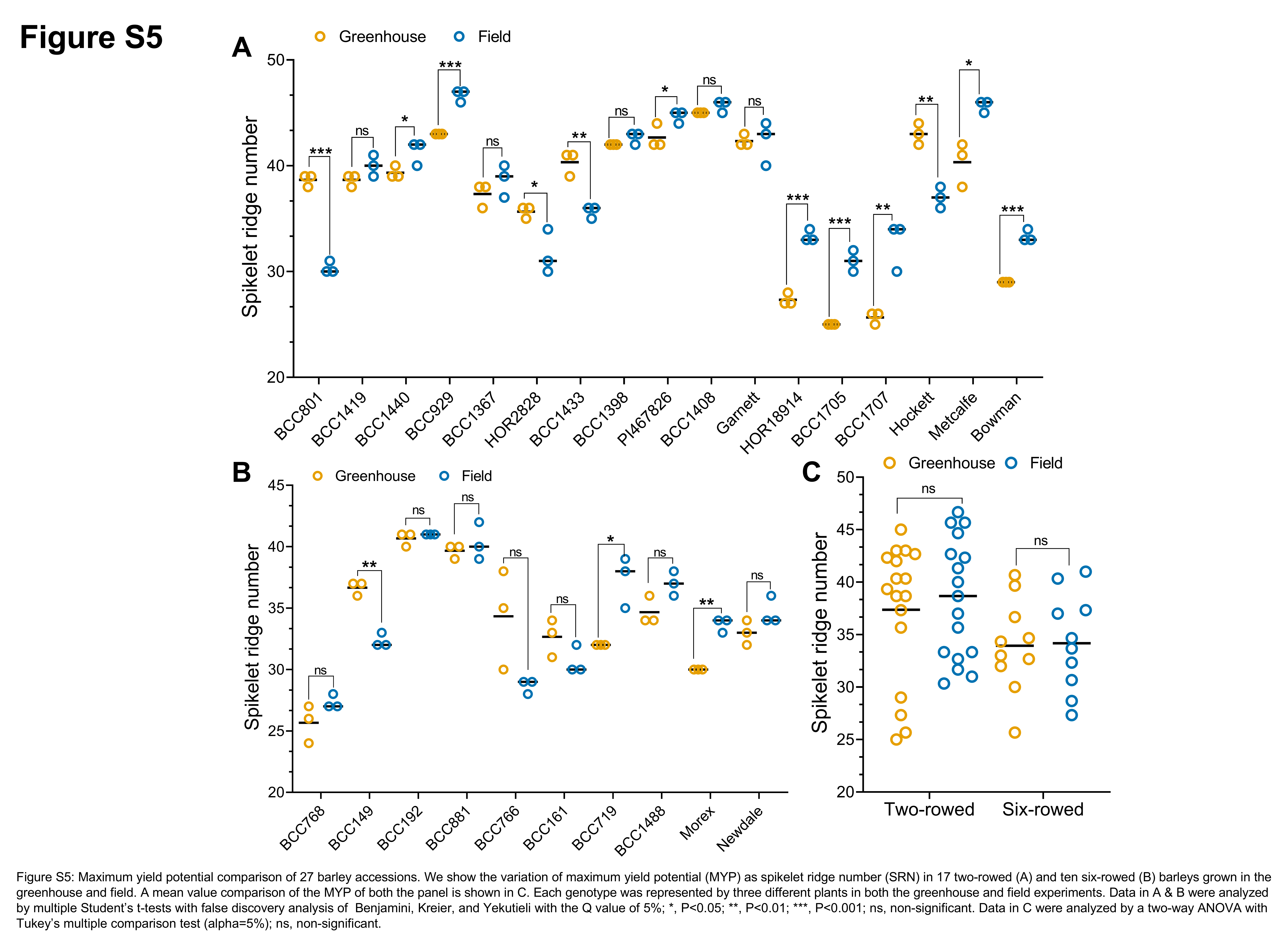

### Figure-S6

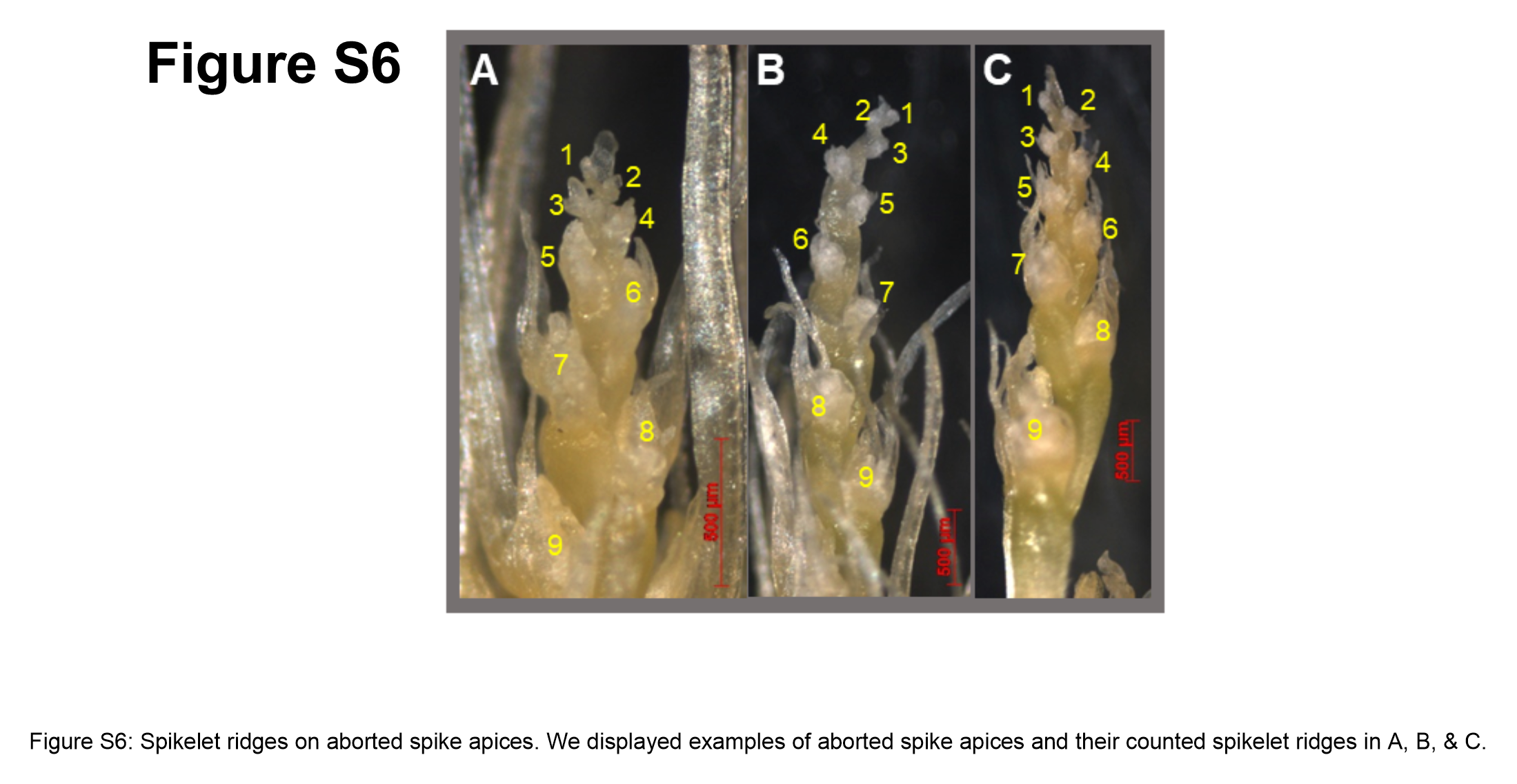
